# Multiple Epigenetic Mechanisms Functionally Cooperate to Silence Expression of Somatostatin Receptor Type 2 in Pancreatic Neuroendocrine Tumors

**DOI:** 10.1101/2025.09.23.677935

**Authors:** James P. Madigan, Stephen G. Andrews, Rivka B. Farrell, Steven D. Forsythe, Rupali Sharma, Michele Ceribelli, Craig J. Thomas, Kimia N. Shamsian, Shinjen Lin, Ken Chih-Chien Cheng, Samira M. Sadowski

## Abstract

Pancreatic neuroendocrine tumors (PNETs) are a rare and understudied set of cancers, with increasing incidence. Neuroendocrine tumors are unique in the fact that they express high levels of the somatostatin receptor type 2 (SSTR2), which represents a target for both tumor imaging and therapeutics. PNET grade inversely correlates with SSTR2 tumor staining and higher tumor grade is associated with poor patient prognosis. With no known mutations, SSTR2 expression is believed to be lost through aberrant epigenetic mechanisms. Enhanced knowledge of the epigenetic biology and players controlling SSTR2 expression may allow for identification of novel PNET imaging and treatment modalities. Through in-depth studies, we found that the specific *de novo* DNA methyltransferase (DNMT), DNMT3B, is responsible for *SSTR2* gene CpG methylation and silencing. Using DNMT3B as a starting point, along with the concept of functional crosstalk between various epigenetic mechanisms, we further discovered that Polycomb Repressor Complexes 1 and 2 (PRC1 and PRC2) play important roles in silencing SSTR2. Moreover, we found several histone lysine demethylases, enzymes that remove activating histone H3K4 methylation marks, to be critical for silencing expression of SSTR2. We additionally identified several chromatin remodeling enzymes/complexes as cellular factors that negatively regulate SSTR2 expression. Finally, using the HiBiT luminescent reporter system, we exploited functional chemo-genomic screens to further expand our knowledge of SSTR2 epigenetic control. These screens both reinforced several of our initial findings and helped to identify additional silencing mechanism potentially regulating SSTR2 expression. A commonality in our findings point to the presence, or necessity, of Class I HDACs in nearly all the epigenetic silencing mechanisms characterized. Overall, our work demonstrates that *SSTR2* gene expression is likely silenced through various dynamic and interconnected epigenetic events, resulting in a compacted, transcriptionally repressed chromatin environment. Our study offers novel potential therapeutic targets and combinations to best increase expression of SSTR2, which are currently being tested in pre-clinical studies from our group, with the goal of future clinical trials aimed at increasing SSTR2 expression in high-grade, SSTR2-low NET patients.

## INTRODUCTION

Pancreatic neuroendocrine neoplasms (PNENs) are a rare set of tumors derived from neuroendocrine cells within the islets of the pancreas and comprising of either pancreatic neuroendocrine tumors (PNETs) or pancreatic neuroendocrine carcinomas (PNECs), separate tumor entities. The World Health Organization has established a classification system to stratify PNENs based on both Ki67 proliferative index and histological differentiation [1]. PNETs are classified under three separate grades: Grade 1 (G1, well differentiated (WD), ki67 < 3%), Grade 2 (G2, WD, ki67 between 3-20%) and Grade 3 (G3, WD, ki67 >20%). PNECs are G3, poorly differentiated and are identified by their aggressiveness and unique mutational spectrum. PNETs are additionally classified as either functional or non-functional tumors, depending on the release of excessive hormones from these tumors. Furthermore, PNETs are most often sporadic, but roughly 10% of cases are linked to specific hereditary syndromes [2]. Due to low tumor mutational burden, the lack of primary oncogenic drivers and the presence of tumor heterogeneity, there has been a dearth of suitable therapeutic strategies to target PNETs.

Unique to PNETs, and NETs in general, is their cell surface expression of Somatostatin Receptor Type 2 (SSTR2). SSTR2 is a member of the SSTR family of G protein coupled receptors, which can be activated by the neuropeptide, somatostatin, resulting in inhibition of both hormone secretion and cell proliferation, along with increased levels of apoptosis [3, 4]. Previous work from our group, at the NCI, led to the FDA-approval of the PET-imaging agent, ^68^Ga-DOTATATE, a high-affinity ligand that binds to SSTR2 [5, 6]. SSTR2 expression on NETs has also led to the ability to therapeutically target tumors via peptide receptor radionuclide therapy (PRRT) using ^177^Lu-DOTATATE [7]. Unfortunately, PRRT is often not effective in high-grade PNET patients, due to loss or decreased expression of SSTR2.

Epigenetics is the study of how the environment and outside factors may alter gene expression, without direct changes to the DNA sequence. Each cell within the body contains the same genetic information, via DNA sequence, yet separate cell types need to phenotypically differentiate into disparate cell types for functional organs and tissues. Epigenetics regulates gene expression through altering the availability of transcription factors acting toward gene targets. The main epigenetic processes include DNA methylation, histone protein post-translational modifications, physical remodeling of chromatin and non-coding RNA activities [8]. In contrast to direct changes to DNA sequences, which are permanent, epigenetic changes are plastic in nature, with the possibility to reverse gene expression alterations. Epigenetic changes, and the resultant gene expression alterations, are facilitated by various proteins and protein complexes that act as either readers, writers or erasers of specific epigenetic modifications [9]. A main feature of epigenetics is the functional crosstalk and physical interaction of various epigenetic proteins, complexes and modifications. This functional crosstalk culminates in either open or closed chromatin structure, resulting in either transcriptionally permissive or non-permissive environments, respectively [10]. Epigenetic changes are needed for normal development, but alterations to normal epigenetic processes can lead to certain genetic disorders and various cancer types [11].

With no known detrimental mutations affecting the *SSTR2* gene, it is believed that loss of SSTR2 expression in NETs is due mainly to epigenetic changes, culminating in compacted chromatin and a transcriptionally non-permissive environment. Until recently, only DNA CpG methylation and histone protein deacetylation have been directly tied to loss of *SSTR2* gene expression. Thus our current work focused on delving deeper into known SSTR2-specific epigenetic silencing mechanisms, with the goal of further clarifying these mechanisms and to identify additional processes responsible for loss of SSTR2 expression. Our results reveal various repressive epigenetic complexes responsible for silencing of SSTR2 expression. A commonality between these repressive multi-subunit complexes is the presence and functional necessity of Class I HDACs, and this fact may play a large part in the efficacy of HDACi in re-expression of SSTR2. Our work represents the most comprehensive analysis of epigenetic-based SSTR2 expression control, to date, and offers possible new therapeutic targets to increase expression of SSTR2, for PNET patient clinical benefit.

## MATERIALS AND METHODS

### Antibodies and reagents

Primary antibodies utilized were: SSTR2 (BosterBio, M01689), DNMT1 (Epigentek, A-1700), DNMT3A (Santa Cruz Biotechnology, sc-365769), DNMT3B (Proteintech, 26971-1-AP), LSH/HELLS (Proteintech, 11955-1-AP) and HiBiT (Promega, N7200). The following antibodies were all obtained from Cell Signaling Technology: GAPDH (#2118), G9a (#68851), SETDB1 (#45389), EZH2 (#5246), RING2/RING1B (#5694), JARID1A/KDM5A (#3876), LSD1 (#2184), RCOR1 (#14567), CHD4 (#12011). HRP-conjugated secondary antibodies were also from Cell Signaling Technology: anti-rabbit IgG (#7074) and anti-mouse IgG (#7076). Therapeutics Decitabine, Tazemetostat (EPZ-6438), Entinostat and Iadademstat (ORY-1001) were purchased from SelleckChem.

### Cell culture

293T cells (American Type Culture Collection, Manassas, VA) were maintained in Dulbecco’s Modified Eagle Media (DMEM), supplemented with 10% fetal bovine serum (FBS) and penicillin-streptomycin. BON-1 cells [12], provided by M. R. Hellmich (University of Texas Medical Branch at Galveston, Galveston, TX), were maintained in DMEM/F-12 media, supplemented with 10% FBS and penicillin-streptomycin. QGP-1 cells [13], purchased from the Japanese Collection of Research Bioresources cell bank, were maintained in RPMI media, supplemented with 10% FBS and penicillin-streptomycin. NT-3 cells [14], provided by Jorg Schrader (University Medical Center Hamburg, Germany), were maintained in RPMI media, supplemented with 10% FBS, penicillin-streptomycin, human FGF-basic and EGF (Peprotech). All cell culture reagents were purchased from Gibco/Life Technologies.

### Western Blot Analysis

After experimental treatments, cells were washed with PBS and lysed with either GPCR Extraction and Stabilization Reagent buffer (#A43436, Thermo Fisher Scientific), or RIPA buffer (Pierce), supplemented with Protease Inhibitor Cocktail (P8340, Millipore-Sigma), rotated at 4°C, sonicated and centrifuged to remove insoluble material. Protein concentrations were determined using a Micro BCA Protein Assay Kit (#23235, Thermo Fisher Scientific). 2× Laemmli sample buffer was added to equivalent amounts of cellular lysates, which were then heated at 95 °C for 5 min and resolved by SDS-PAGE on Novex 4–12% Bis-Tris gels (Invitrogen) and transferred onto Immobilon polyvinylidene difluoride membrane (Millipore).

Membranes were blocked in 5% (w/v) nonfat dried skimmed milk powder in TBS-Tween 20 (blocking buffer) (20 mm Tris/HCl, pH 7.6, 137 mm NaCl, and 0.2% Tween 20) and probed with appropriate primary antibodies followed by anti-mouse or anti-rabbit IgG-horseradish peroxidase–conjugated secondary antibody (Cell Signaling Technology). Membranes were washed in TBS-Tween 20 and incubated in Immobilon Western Chemiluminescence substrate (Millipore). Band intensities were quantified using Image J software (National Institutes of Health) and normalized to loading control protein expression.

### Targeted NextGen Bisulfite Sequencing (tNGBS)

Targeted NextGen Bisulfite Sequencing was performed by EpigenDx, Inc. (Hopkinton, MA).

#### Assay Design

Each regulatory element of a requested gene was carefully evaluated before beginning the process of assay design. Gene sequences containing the target of interest were acquired from the Ensembl genome browser and annotated. The target sequences were re-evaluated against the UCSC genome browser for repeat sequences including LINE, SINE, and LTR elements. Sequences containing repetitive elements, low sequence complexity, high thymidine content, and high CpG density were excluded from the *in silico* design process.

#### Sample Digestion

Cell pellet samples were lysed based on the total cell count per sample and total volume received using M-digestion Buffer (ZymoResearch; Irvine, CA; cat# D5021-9) and 5-10µL of protease K (20mg/mL), with a final M-digestion concentration of 2X. The samples were incubated at 65°C for a minimum of 2 hours.

#### Bisulfite Modification

20µL of the supernatant from the sample extracts were bisulfite modified using the EZ-96 DNA Methylation-Direct Kit™ (ZymoResearch; Irvine, CA; cat# D5023) as per the manufacturer’s protocol with minor modification. The bisulfite modified DNA samples were eluted using M-elution buffer in 46µL.

#### Multiplex PCR

i. All bisulfite modified DNA samples were amplified using separate multiplex or simplex PCRs. PCRs included 0.5 units of HotStarTaq (Qiagen; Hilden, Germany; cat# 203205), 0.2µM primers, and 3µL of bisulfite-treated DNA in a 20µL reaction. All PCR products were verified using the Qiagen QIAxcel Advanced System (v1.0.6). Prior to library preparation, PCR products from the same sample were pooled and then purified using the QIAquick PCR Purification Kit columns or plates.
ii. Recommended PCR cycling conditions: 95°C 15 min; 45 x (95°C 30s; **Ta**°C 30 s; 68°C 30 s); 68°C 5 min; 4°C ∞
iii. Samples were run alongside established reference DNA samples with a range of methylation. They were created by mixing high- and low-methylated DNA to obtain samples with 0, 50, and 100% methylation. The high-methylated DNA is *in vitro* enzymatically methylated genomic DNA with >85% methylation. The low-methylated DNA is chemically and enzymatically treated with <5% methylation. They are first tested on numerous gene-specific and global methylation assays using pyrosequencing.

#### Library Preparation and Sequencing

Libraries were prepared using a custom Library Preparation method created by EpigenDx. Next, library molecules were purified using Agencourt AMPure XP beads (Beckman Coulter; Brea, CA; cat# A63882). Barcoded samples were then pooled in an equimolar fashion before template preparation and enrichment were performed on the Ion Chef™ system using Ion 520™ & Ion 530™ ExT Chef reagents (Thermo Fisher; Waltham, MA; cat# A30670). Following this, enriched, template-positive library molecules were sequenced on the Ion S5™ sequencer using an Ion 530™ sequencing chip.

#### Data Analysis

FASTQ files from the Ion Torrent S5 server were aligned to a local reference database using the open-source Bismark Bisulfite Read Mapper program (v0.12.2) with the Bowtie2 alignment algorithm (v2.2.3). Methylation levels were calculated in Bismark by dividing the number of methylated reads by the total number of reads. An R-squared value (RSQ) was previously calculated from controls set at known methylation levels to test for PCR bias.

### shRNA-mediated stable knockdown

To stably knock down cellular expression of specific epigenetic players, the pLKO.1-puro lentivirus plasmid-based MISSION shRNA system was used. Human-specific shRNA clones were initially selected for highest in-house experimental knockdown (Millipore-Sigma, Broad Institute) and final selection based on validation of superior knockdown from results of independent research sources. The following specific shRNA clones were purchased from Millipore-Sigma: DNMT1 (TRCN0000021893), DNMT3A (TRCN0000035758), DNMT3B (TRCN0000035686), EHMT2 (G9A) TRCN0000115670, SETDB1 (TRCN0000148112), EZH2 (TRCN0000040074), RING2 (RNF2) (TRCN0000033697), KDM5A (JARID1A) (TRCN0000329872), KDM1A (LSD1) (TRCN0000046071), HELLS (LSH) (TRCN0000273217), RCOR1 (CoREST1) (TRCN0000128570), CHD4 (TRCN0000021363). Additionally, a non-targeting shRNA plasmid (NT-shRNA) that targets no known human sequence was utilized as a control [15]. A primer containing the target sequence (CTGGTTACGAAGCGAATCCTT) along with a stem loop followed by the reverse target sequence was annealed to a complementary primer and inserted into the EcoRI and AgeI sites of pLKO.1-puro (Addgene number 10878). Lentiviral particles were produced via Lipofectamine 2000 (Invitrogen)-mediated triple transfection of 293T cells with the respective pLKO.1-puro shRNA plasmids along with the lentiviral envelope plasmid (pMD2.G, Addgene number 12259) and the lentiviral packaging plasmid (psPAX2, Addgene number 12260). Target cells were transduced with shRNA containing lentiviral particles in the presence of 8 μg/ml Polybrene, and stable cells were selected using 2 μg/ml puromycin.

### Quantitative high-throughput single-agent screening (qHTS)

High-throughput single agent drug screenings were performed as previously described [16]. Briefly, BON-1 SSTR2-HiBiT cells (clone #2) were seeded in 5 μL of growth media using a Multidrop Combi dispenser (ThermoFisher) into 1536-well white polystyrene tissue culture-treated plates (Aurora) at a density of 2000cells/well. 23 nL of MIPE 6.0 compounds were added to individual wells (11 doses tested for each compound in separate wells, for a total of 22x1536-well plates for each cell-line/readout) via a 1536 pin-tool. Plates were incubated for 24h at standard incubator conditions covered by a stainless steel gasketed lid to prevent evaporation.

To measure the activity of the HiBiT-based reporter, 4 μL of Nano-Glo® HiBiT Lytic Detection Reagent (Promega) were added to each well and plates were incubated at room temperature for 15 min with the stainless-steel lid in place. Luminescence readings were taken using a Viewlux imager (PerkinElmer) with a 60’’ exposure time and a 2X binning per plate. Relative reporter activity was assessed with respect to DMSO treated wells.

Viability was assessed in parallel in a separate set of MIPE 6.0 plates. Briefly, following 24h of drug exposure, 3 μL of Cell-Titer Glo® (Promega) were added to each well. Plates were incubated at room temperature for 15 min with the stainless-steel lid in place. Luminescence readings were taken using a Viewlux imager (PerkinElmer) with a 2’’ exposure time and 1X binning per plate. Relative viability was assessed with respect to DMSO treated wells (column #4 in each plate). The criteria we used to batch-classify dose-response curves based on the quality of curve fit had been previously described [17].

### Quantitative pairwise combination screening

Pairwise drug-combination screenings were performed as previously described [18, 19]. Briefly, 10 nL of compounds were acoustically dispensed into 1536-well white polystyrene tissue culture-treated plates with an Echo 550 acoustic liquid handler (Labcyte). Cells were then added to compound-containing plates at a density of 2000-cells/well in 5 μL of medium. A 10-point custom concentration range, with constant 1:2 dilution, was used for all the drug combination pairs assessed in 10 × 10 matrix format. Synergistic effects on reporter activity and/or viability were assessed by 24h post drug treatment, as described above for single agent qHTS.

### Statistical Analysis

Western blot band densitometry was performed with ImageJ software (National Institutes of Health). SSTR2 protein band intensity was compared between control and drug-treated samples and normalized to loading control band intensity (GAPDH). Data analysis was performed using an unpaired Student’s *t*-test using Prism software (Graphpad). *P* values < 0.05 were considered statistically significant.

## RESULTS

### Control of SSTR2 gene expression via de novo DNA CpG methylation

Work from our lab, and others, has previously identified broad-based epigenetic silencing mechanisms, such as DNA CpG methylation and histone deacetylation, in negatively controlling SSTR2 expression [20, 21]. Regarding DNA CpG methylation, studies to date have used non-selective DNMT inhibitors, such as Azacitidine and Decitabine, to increase expression of SSTR2. As seen in **Fig 1A**, treatment with Decitabine resulted in over 4.5- and 6-fold increases in SSTR2 protein levels in BON-1 and QGP-1 PNET cell lines, respectively, compared to vehicle control. To determine precisely which DNMT(s) are responsible for methylation of CpG sites of *SSTR2* gene regulatory elements, stable knockdown using pre-validated shRNAs targeting human DNMT1, DNMT3A and DNMT3B were employed, along with a control shRNA targeting no known human gene (NT-shRNA). We confirmed high level knockdown of each DNMT in the BON-1 cell line (**Fig. 1B**). While stable knockdown of both DNMT1 and DNMT3A had no effect on SSTR2 protein levels, stable knockdown of DNMT3B resulted in increased total SSTR2 protein expression in both BON-1 and QGP-1 cells (**Fig. 1C**). Using next-generation bisulfite sequencing analysis, we demonstrated that stable knockdown of DNMT3B significantly decreased DNA methylation at specific CpG sites around the *SSTR2* promoter region (**Fig. 1D**). Finally, using a PNET cell line with high level expression of SSTR2, NT-3 [14], compared to SSTR2-low BON-1 and QGP-1 cells, we demonstrated a correlation between high levels of SSTR2 expression and decreased SSTR2 promoter CpG methylation, downstream of the transcription start site (**Fig. 1E**). These results reconfirm that DNA CpG methylation is an important epigenetic mechanism involved in *SSTR2* gene silencing, and that the *de novo* DNMT, DNMT3B, is responsible for this silencing in BON-1 and QGP-1 cell lines.

**Figure 1:**
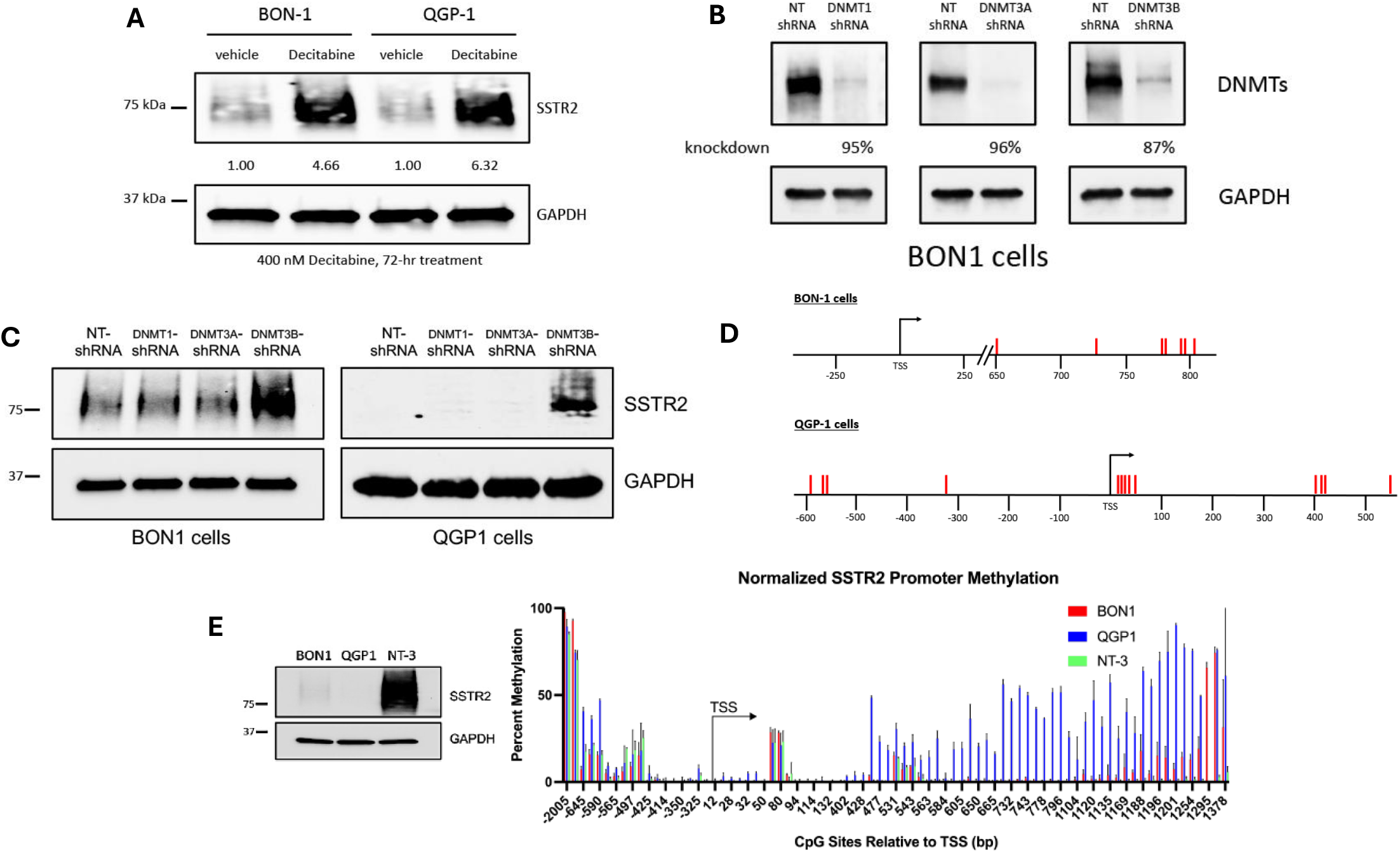
Role and characterization of SSTR2 DNA methylation. **(A)** Immunoblot analysis of BON-1 and QGP-1 cells treated with the DNMT inhibitor, Decitabine (400 nM for 72-hrs) shows increased expression of total SSTR2 protein. **(B)** Immunoblot analysis confirms stable knockdown of DNMTs in BON-1 cells using pre-validated shRNA targeting DNMT1, DNMT3A and DNMT3B. Specific percent stable knockdown of each DNMT is noted (similar knockdown results in QGP-1 cells). **(C)** Immunoblot analysis reveals that stable knockdown of DNMT3B in both BON-1 and QGP-1 cells selectively increases expression of total SSTR2 protein. **(D)** Schematic of the human *SSTR2* gene promoter element and CpG sites (red lines) having significantly decreased methylation levels (P < 0.05, at the least) after stable knockdown of DNMT3B in BON-1 and QGP-1 cells, compared to cells expressing control non-targeting shRNA (n=3). **(E)** Total protein levels in BON-1, QGP-1 and NT-3 cells correlates with *SSTR2* gene promoter CpG methylation levels (n=3).

### Polycomb repressor complexes in control of SSTR2 expression

Applying our discovery of DNMT3B in controlling SSTR2 expression, we used the fact of interconnectedness between various epigenetic mechanisms to help uncover additional epigenetic processes responsible for SSTR2 silencing. Importantly, evidence suggests of crosstalk between CpG DNA methylation and histone-based epigenetic marks, specifically histone lysine methylation [22]. Inhibitory methylation marks at histone H3K9 and direct binding to H3K9-specific lysine methyltransferases have been demonstrated to control *de novo* methylation of heterochromatin by both DNMT3A and B [23]. In BON-1 cells (**Fig. 2A**), stable shRNA-mediated knockdown of two separate H3K9 methyltransferases (G9a and SETDB1) did not lead to SSTR2 re-expression, with stable knockdown of DNMT3B as a positive control. Similar results were seen in the QGP-1 cell line. These results suggest that H3K9 methylation-dependent transcriptional repression mechanisms may not be pertinent to silencing of SSTR2 expression. An additional inhibitory histone methylation mark, H3K27me3, laid down by the histone methyltransferases, EZH1/2, of the Polycomb Repressor Complex 2 (PRC2), has also been shown to have functional crosstalk with DNA methylation. While the interplay between H3K27me3 and DNA CpG methylation is complex [22], it has been demonstrated that the PRC2 enzymatic subunit, EZH2, directly controls DNA methylation by binding to DNMTs and guiding them to polycomb group repressed genes [24]. A search for possible connections between PRC2 complex signaling and SSTR2 expression found that multiple “GPCR genes” are subject to PRC2-mediated H3K27me3 modifications [25], and independent ChIP-seq studies have found the *SSTR2* gene to be bound by PRC2 complexes and contain the H3K27me3 repressive mark [26, 27]. Importantly, we demonstrate that stable shRNA-mediated knockdown of EZH2, in both BON-1 and QGP-1 cells, led to increased expression of total SSTR2 protein (**Fig. 2B**). Additionally, treatment with the EZH2-specific inhibitor, Tazemetostat [28], led to statistically significant increased expression of total SSTR2 protein in both BON-1 and QGP-1 cells (**Fig. 2C**).

**Figure 2:**
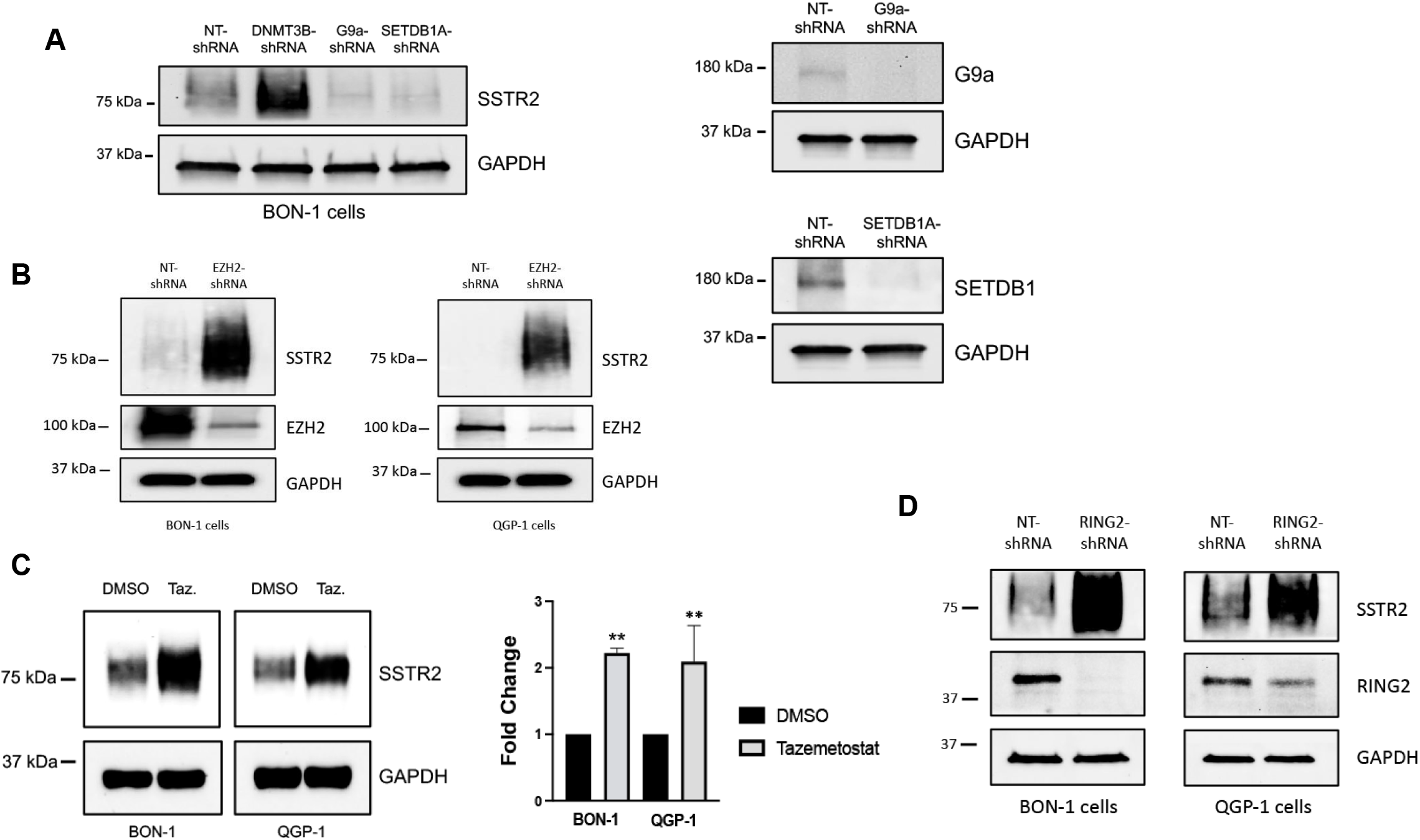
A role for Polycomb Repressor Complexes 1 and 2 (PRC1 and 2) in controlling SSTR2 expression. **(A)** Immunoblot analysis demonstrates that stable knockdown of H3K9 histone methyltransferases, G9A and SETDB1, do not affect total SSTR2 protein expression. **(B)** Immunoblot analysis reveals that stable knockdown of the H3K27 histone methyltransferase, EZH2, increases expression of total SSTR2 protein in both BON-1 and QGP-1 cells. **(C)** Immunoblot analysis of both BON-1 and QGP-1 cells treated with the EZH2-specific inhibitor, Tazemetostat (1 uM for 96-hrs). Statistical analysis confirms significantly increased expression of SSTR2 protein expression after treatment with Tazemetostat. Data are expressed as mean ± SEM, **P < 0.01 (n=3). **(D)** Immunoblot analysis demonstrates that stable knockdown of RING2, a catalytic subunit of PRC1, increases expression of total SSTR2 protein.

Polycomb Repressor Complex 1 (PRC1) is a PRC2-related polycomb repressor-based epigenetic system and is responsible for writing the repressive H2AK119Ub mark [29]. Previous studies have demonstrated intimate feedforward and feedback signaling between PRC1 and PRC2 complexes, and the marks they lay down, as important for optimal silencing of polycomb target genes [30]. As shown in **Fig. 2D**, stable shRNA-mediated knockdown of RING2, a catalytic subunit of PRC1 [31], led to increased expression of total SSTR2 protein in both BON-1 and QGP-1 cell lines. These results suggest that *SSTR2* gene expression is negatively regulated by PRC1/2 epigenetic silencing mechanisms and may provide future valuable therapeutic targets to increase expression of SSTR2 in SSTR2-low NET patients.

### Specific histone lysine demethylases control SSTR2 expression

The concept of bivalency suggests that the chromatin of certain gene promoters are decorated by both activating and inhibitory histone methylation marks, in a “poised state”, and bivalent genes can therefore rapidly transfer from active to repressive transcriptional states [32]. Bivalent promoters often contain activating H3K4me2,3 and inhibitory H3K27me3 methylation marks. After uncovering a role for PRC1/2-mediated repression of SSTR2 expression, we sought a role for possible demethylases that may function to silence *SSTR2* gene expression through removal of activating marks at H3K4. The JARID1 (KDM5) family of H3K4 demethylases (JARID1A, B, C and D) have been shown to play roles in a variety of cancer types [33]. While JARID1B is the best characterized family member, JARID1A has been shown to play a functional role in PNETs [34]. Stable shRNA-mediated knockdown of JARID1A (**Fig. 3A**), resulted in increased levels of total SSTR2 protein in both BON-1 and QGP-1 cell lines.

**Figure 3:**
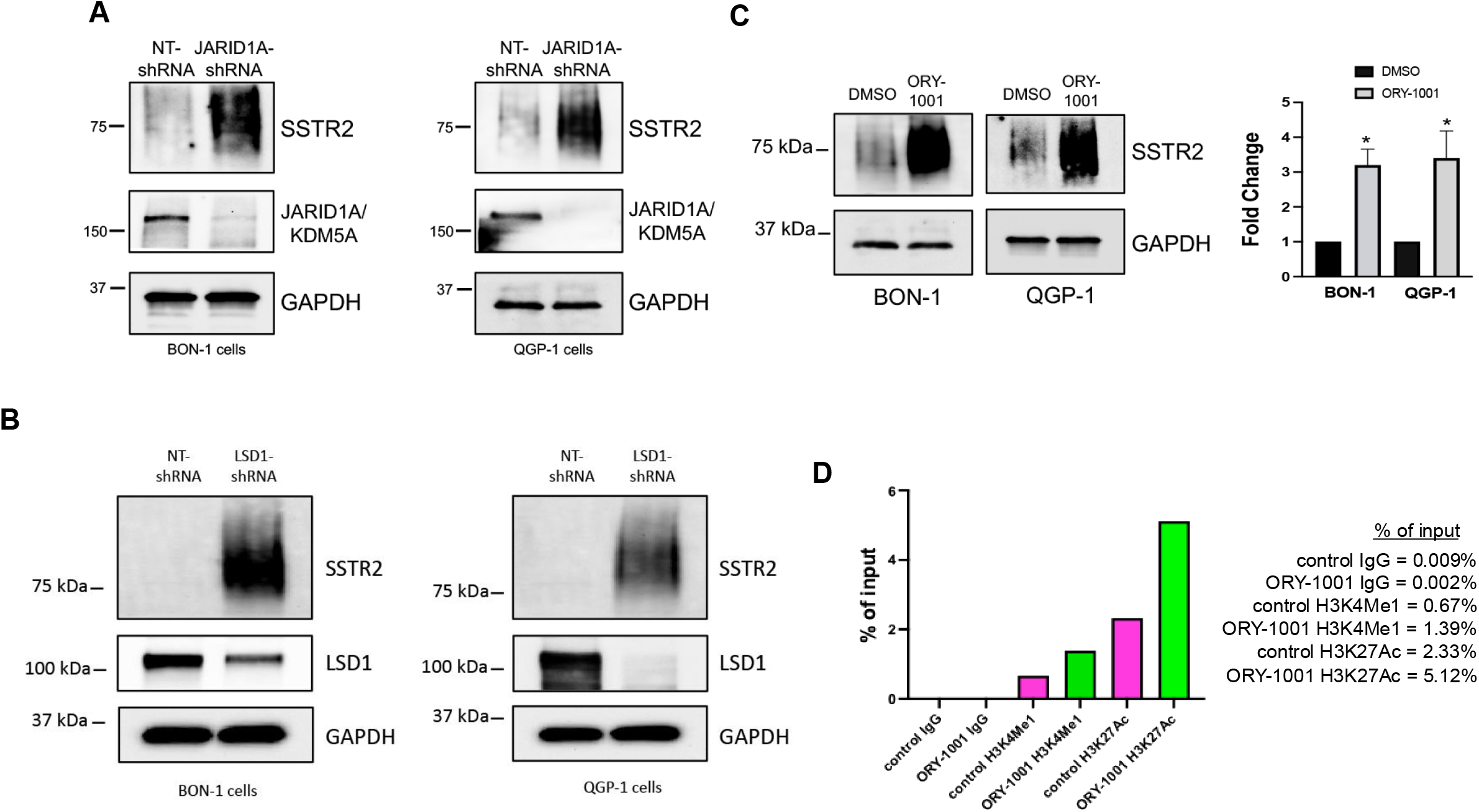
The histone lysine demethylase, LSD1, controls expression of SSTR2 and regulates enhancer element histone modifications. **(A)** Immunoblot analysis demonstrates that stable knockdown of the histone H3K4 lysine demethylases, JARID1A, increases expression of total SSTR2 protein in both BON-1 and QGP-1 cells. **(B)** Immunoblot analysis reveals that stable knockdown of LSD1, in both BON-1 and QGP-1 cells, increases expression of total SSTR2 protein. **(C)** Immunoblot analysis of both BON-1 and QGP-1 cells treated with the LSD1-specific inhibitor, ORY-1001 (1 uM for 72-hrs). Statistical analysis confirms significantly increased expression of SSTR2 protein after ORY-1001 treatment. Data are expressed as mean ± SEM, *P < 0.05 (n=3) **(D)** ChIP analysis demonstrates that treatment with ORY-1001 selectively increases activating marks, H3K4me1 and H3K27Ac, within the SSTR2 enhancer.

The first histone lysine demethylase discovered was LSD1 (KDMA1A) [35] and has been demonstrated to remove H3K4me1,2 activating marks, often found at gene enhancer elements. Stable shRNA-mediated knockdown of LSD1, in both BON-1 and QGP-1 cell lines, resulted in increased levels of total SSTR2 protein (**Fig. 3B**). Additionally, treatment of both BON-1 and QGP-1 cell lines with the LSD1-specific inhibitor, Iadademstat (ORY-1001) [36], also led to statistically significant increased expression of total SSTR2 protein in both BON-1 and QGP-1 cells (**Fig. 3C**). Importantly, ChIP analysis of the *SSTR2* gene enhancer element showed that treatment of QGP-1 cells with Iadademstat resulted in increased levels of two activating marks, H3K4me1 and H3K27Ac, often found at enhancers of transcribed genes (**Fig. 3D**). Our results suggest that multiple histone lysine demethylases play a role in silencing of SSTR2 expression and may provide additional future therapeutic targets to increase SSTR2 levels in patients.

### Chromatin remodeling factors affect SSTR2 expression

LSD1 does not act alone but is a component of several distinct repressor complexes [37]. The first complex found to contain LSD1 was CoREST, a chromatin-modifying corepressor complex that regulates gene expression [38]. Stable knockdown of the CoREST member, RCOR1 [39], did not increase total SSTR2 protein levels in either BON-1 or QGP-1 cells (**Fig. 4A**). An additional repressor complex shown to contain LSD1 is the Nucleosome Remodeling Deacetylase (NuRD) complex [40]. Unlike our results with stable knockdown of RCOR1, stable knockdown of the core nucleosome remodeling component of the NuRD complex, CHD4 [41], increased levels of total SSTR2 protein in both BON-1 and QGP-1 cells (**Fig. 4B**). As the NuRD complex contains chromatin remodeling activity through CHD4 ATPase-dependent activity, we additionally tested a possible role for the SNF2 family helicase, Lymphoid-Specific Helicase (LSH), in control of SSTR2 expression. As seen in **Fig. 4C**, stable knockdown of LSH in both BON-1 and QGP-1 cells increased levels of total SSTR2 protein. Beyond ATP-dependent chromatin remodeling activity, LSH has been shown to affect *de novo* DNA methylation, through control of DNMT3A/B activity [42] and may represents an additional example of epigenetic crosstalk in control of SSTR2 expression. Our results suggest that in addition to DNA methylation and histone modifications, specific chromatin remodeling factors, often found in large multi-subunit complexes, likely play important roles in regulating expression of SSTR2.

**Figure 4:**
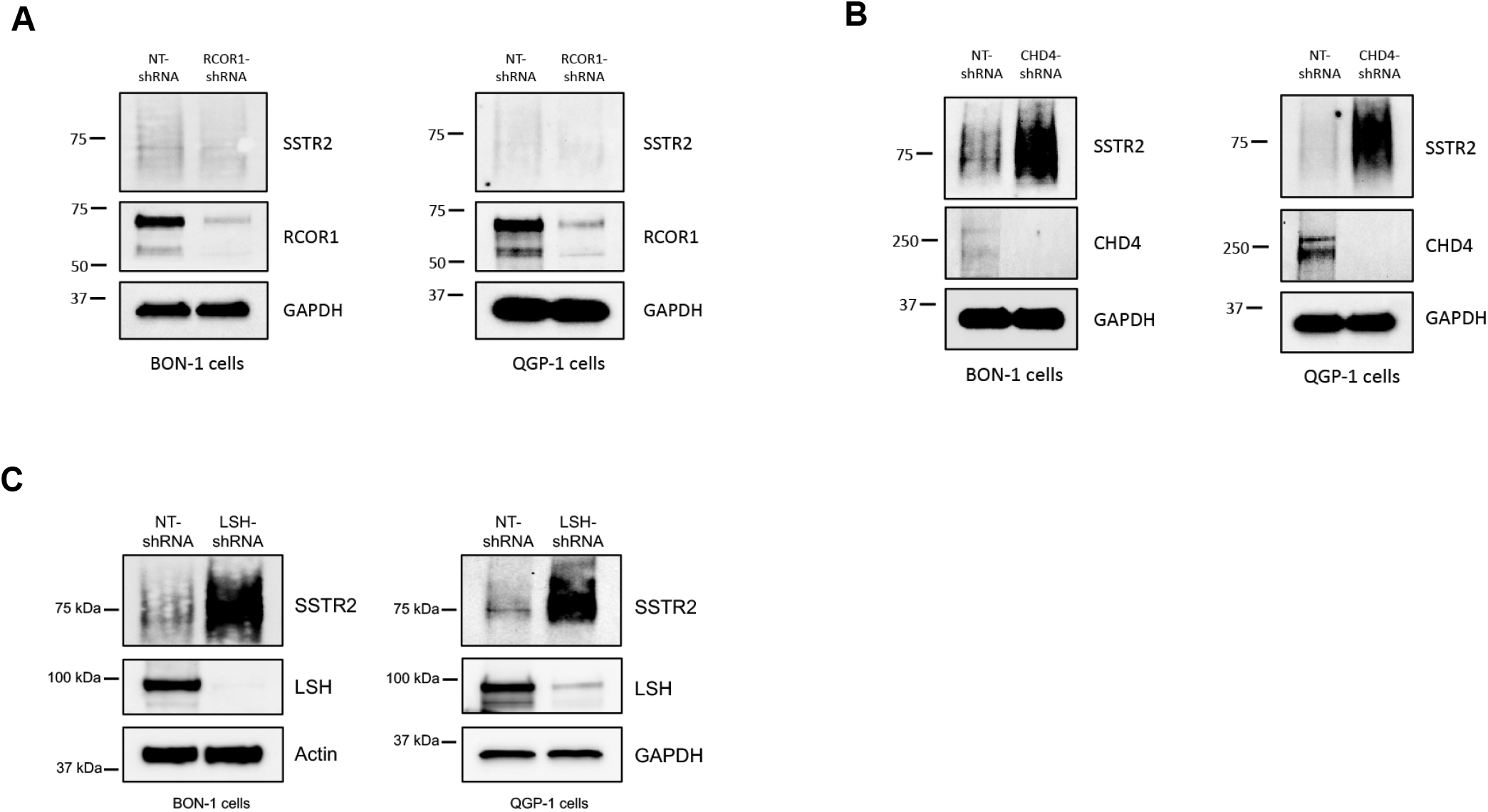
Role of chromatin remodeling factors in controlling SSTR2 expression. **(A)** Immunoblot analysis demonstrates that stable knockdown of the CoREST subunit, RCOR1, in both BON-1 and QGP-1 cells does not affect total SSTR2 protein expression. **(B)** Immunoblot analysis reveals that stable knockdown of the essential NuRD complex factor, CHD4, increases total SSTR2 protein expression in both BON-1 and QGP-1 cells. **(C)** Immunoblot analysis shows that stable knockdown of the chromatin-remodeling protein, LSH, increases total SSTR2 protein in both BON-1 and QGP-1 cells.

### Functional genomic and drug screening

To identify additional factors controlling expression of SSTR2 and further validate our findings, the HiBiT luminescent reporter system was employed [43]. Through CRISPR gene editing technology, the 11 amino acid HiBiT tag was inserted into the extreme C-terminal coding sequence of the *SSTR2* gene, just upstream of the stop codon. A clone of the BON-1 PNET cell line, with one *SSTR2* allele knocked-in with the HiBiT reporter tag, was used for semi-biased functional genomic and chemical screening (**Fig. 5A**). Using a Dharmacon epigenetic-focused siRNA library to knockdown specific epigenetic-related genes, we confirmed some of our initial findings. For example, siRNA-mediated knock down of members of the PRC2 complex (EZH2, SUZ12 and EED) along with the co-repressor, CDYL, which promotes a feedback loop for PRC2 signaling [44], members of the NuRD complex (CHD3 and MBD3) and LSD (KDM1A) itself, all increased HiBiT signal. We also identified additional possible regulators of SSTR2 expression. Some of the most promising targets whose specific knockdown led to increased HiBiT luminescent signal were: **CDK5R1** (p35, neuron-specific activator of CDK5), **CECR2** (histone acetyl-lysine reader), **ANKRD1** (ankyrin repeat protein 1), **VDR** (Vitamin D Receptor), **DMAP1** (DNA methyltransferase 1 associated factor), **SKI** (SKI proto-oncogene), **DIP2B** (disco interacting protein 2 homolog B; works in conjunction with DMAP1), **TLE5** (TLE family member 5, transcriptional modulator), **PWWP2A** and **PWWP2B** (PWWP domain containing 2A and B; members of an alternative NuRD repressor complex), **JUND** (JunD proto-oncogene; AP-1 transcription factor subunit) and **HMG20B** (high mobility group 20B).

**Figure 5:**
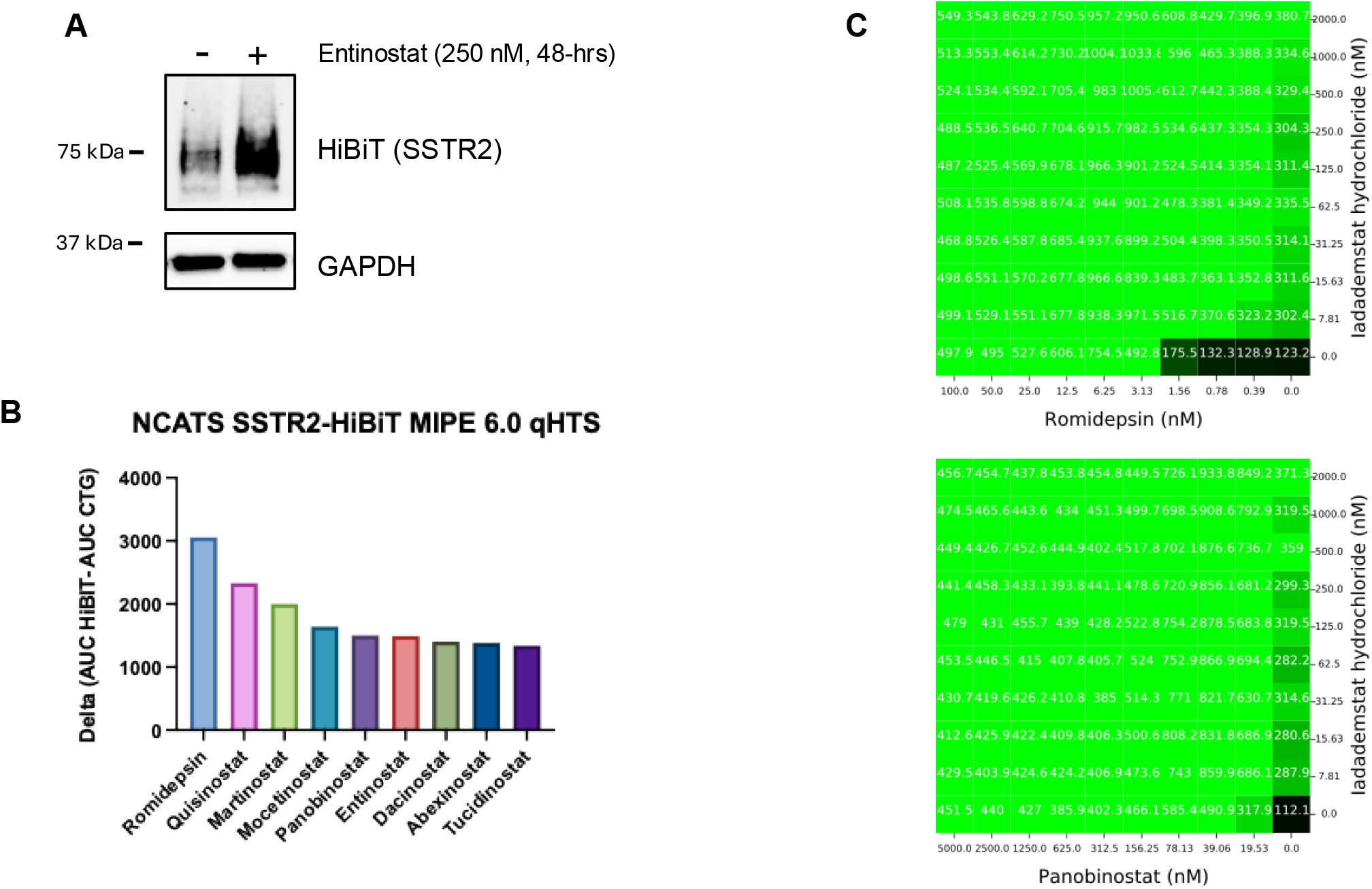
SSTR2-HiBiT-coupled chemo-genomic functional screens. **(A)** Immunoblot analysis demonstrates that treatment with the HDACi, Entinostat (250 nM for 48-hrs), increases expression of total SSTR2-HiBiT protein and confirms that SSTR2-HiBiT expression is dynamically regulated by epigenetic-modifying compounds. **(B)** Quantitative high-throughput screening, using the MIPE 6.0 library, demonstrates 9 out of the top 10 top hits being members of the HDACi family. **(C)** Example combination drug screening of top hit drugs from primary screen. Rhomidepsin + Iadademstat and Panobinostat + Iadademstat. Readout is HiBiT luminescence.

While this functional genomic screen provided deeper insight into the potential epigenetic biology of *SSTR2* gene regulation, we sought further translational relevance, with the goal of performing future clinical trials, focused on SSTR2-based therapeutics against PNETs. Doing quantitative high-throughput screening, using the MIPE 6.0 library (containing 2803 mechanistically annotated drugs), we also sought to discover potential targetable regulators of *SSTR2* gene expression. Nine out of the top 10 (**Fig. 5B**) and 15 out of the top 20 hits were HDACi. Also scoring high was inhibition of LSD1/KDM1A. Inhibition of BMI1 (member of the PRC1 complex) demonstrated high potential, as interfering with the PRC1 complex, in our hands, also increased expression of SSTR2 protein (**Fig. 2**).

Combination drug screening using several of the top hits and classes was also performed, with the rationale for discovery of drug combinations that provide the greatest increase in SSTR2 expression, but also with the lowest drug concentrations, to avoid unnecessary patient toxicity. In line with our previous findings, LSD1 inhibition, using Iadademstat (ORY-1001), provided some of the most impressive additive/synergistic increases in HiBiT-SSTR2 expression, when added with top scoring HDAC inhibitors Rhomidepsin and Panobinostat (**Fig. 5C**). These chemo-genomic screening results have not only provided additional clues on how *SSTR2* gene expression is epigenetically regulated, but also potential therapeutic targets for future SSTR2-focused PNET therapeutics.

## DISCUSSION

Little is known regarding the epigenetic events and players, at the molecular level, which may coordinate to negatively control SSTR2 expression. Distinct epigenetic mechanisms often work in concert with one another through functional crosstalk, and we analyzed these interactions in our studies to uncover novel roles for various epigenetic players and marks in controlling SSTR2 expression. Initially, with the knowledge that DNMT inhibition results in increased expression of SSTR2 [45], we discovered that DNMT3B is key for *SSTR2* gene methylation and silencing. Using DNMT3B as our experimental starting point, we discovered a role for the enzymatic component of the PRC2 complex, EZH2, which lays down the repressive H3K27me3 histone mark [46], in control of SSTR2 expression. Activities of the PRC2 complex, along with the functionally connected PRC1 complex, have been co-opted by cancer, as members of both complexes are found to be overexpressed in numerous cancers and directly correlate with poor patient outcomes [47]. Apropos to our investigation of epigenetic control of SSTR2, both PRC2 [24, 48, 49] and PRC1 [50, 51] serve as recruitment platforms for DNMT3B. Initially, it was believed that DNA methylation and the repressive H3K27me3 mark were mutually exclusive at gene regulatory elements. Contrary to this thought, it has been demonstrated in cancer cells that there is coexistence of both marks at certain gene promoters [52, 53]. Importantly, genes targeted by PRC1/2 complexes are generally associated with high-density CpG promoters [54] and genes that are CpG methylated in cancer are frequently marked by PRC binding and H3K27 methylation early in development [55]. Furthermore, in colon cancer, genes subject to tumor-specific methylation were more likely to be marked by H3K27me3 [56] and 47% of DNMT3B-regulated genes were found to be bound by PRC1/2 [50]. These findings highlight an intimate connection between DNA methylation and polycomb-mediated repression systems. Notably, therapeutic EZH2 inhibition is currently being investigated in PNENs [57]. Our data demonstrates an increase in total SSTR2 protein after both stable knockdown and inhibition of EZH2 and therefore, dual treatment with DNMT and EZH2 inhibitors may represent a potential treatment option to increase expression of SSTR2.

Gene promoters in cancer are often found to be bivalently marked with both activating and inhibitory chromatin marks, and recent work has demonstrated that bivalent chromatin in cancer cells protects genes from *de novo* DNA methylation [58]. We demonstrate here that stable knockdown of JARID1A, a histone lysine demethylase which removes activating H3K4me2,3 marks, increases expression of total SSTR2 protein levels. Interestingly, the PRC2 complex recruits JARID1A to its target genes, and that this interaction is required for PRC2-mediated transcriptional repression during ES cell differentiation [59]. It is therefore plausible that during *SSTR2* gene silencing, histone lysine demethylases, such as JARID1A, remove activating H3K4 histone methylation, in tandem with increased layering of PRC2-dependent inhibitory H3K27 methylation marks.

In additional to JARID1A, we found that both stable knockdown and inhibition of an additional histone lysine demethylase, LSD1, increased levels of total SSTR2 protein. Recent preclinical phase 1 studies have demonstrated promising results using an orally available LSD1 inhibitor in various neoplasms [60, 61]. A separate research group recently demonstrated the potential of targeting LSD1, along with HDAC inhibition, to increase expression of SSTR2 in PNETs [62]. Results from our combination drug screens also suggests that dual HDAC and LSD1 inhibition may be a potential formulation to increase levels of SSTR2 in patients.

LSD1 has been previously shown to functionally interact with several multi-subunit repressor complexes, including the NuRD complex [40]. In our hands, stable knockdown of the core nucleosome remodeling component of NuRD, CHD4, increases expression of total SSTR2 protein. Importantly, the NuRD complex has been shown to functionally interact with several epigenetic players we have found to control SSTR2 protein expression, including JARID1A [63] and PRC2 [64, 65], adding a potential further layer of functional epigenetic crosstalk in control of SSTR2 expression.

Class I HDACs, HDAC1/2, are often associated with multi-subunit repressor complexes, such as the Sin3, NuRD and CoREST complexes [66]. Additionally, Class I HDACs have been found to be intimately connected to both LSD1 [67] and LSH [68] and essential for their biological activity. Likewise, there is experimental evidence demonstrating that Class I HDACs also functionally interact with the PRC2 complex [69, 70]. Furthermore, DNMT3B has been found to bind and crosstalk with Class I HDACs [71]. Thus, nearly all the novel epigenetic mechanisms we have investigated in our current study, along with numerous high scoring hits from our chemo-genomic screens functionally crosstalk or require Class I HDACs. The functional necessity of Class I HDACs may explain the general utility of HDAC inhibition to increase SSTR2 expression demonstrated by several research groups, including our own [20, 72-74].

In summary, our study has expanded our knowledge of the epigenetic biology controlling SSTR2 expression and has uncovered roles for several epigenetic players utilizing various epigenetic mechanisms, including DNA methylation, histone modifications (both writers and erasers) and chromatin remodeling. Our results suggest that SSTR2 silencing likely involves coordinated input from these various epigenetic mechanisms to create a silent, compressed chromatin environment. With multiple layers of epigenetic control likely silencing *SSTR2* gene expression, optimal re-expression of silenced SSTR2 expression may require targeting of two or more separate epigenetic mechanisms. Through chemo-genomic functional screens, our study offers potential therapeutic targets and combinations to best increase expression of SSTR2. These drug combinations are currently being tested in pre-clinical studies from our group, with the goal of future clinical trials aimed at increasing SSTR2 expression in high-grade, SSTR2-low NET patients.

